# Placental lipid handling, growth and inflammatory pathways are modified by a maternal Mediterranean diet

**DOI:** 10.64898/2026.03.23.711314

**Authors:** J Lopez-Tello, L Youssef, R Bermejo-Poza, A Cabeza, J De la Fuente, F Crovetto, E Gratacos, F Crispi, AN Sferruzzi-Perri

**Affiliations:** Department of Physiology, Development, and Neuroscience, The Loke Centre for Trophoblast Research, University of Cambridge, Cambridge, UK; Instituto de Investigaciones Biomédicas Sols-Morreale (IIBm, CSIC-UAM), Madrid, Spain; Department of Physiology, Faculty of Medicine, Autonomous University of Madrid, Madrid, Spain; BCNatal Fetal Medicine Research Center (Hospital Clínic and Hospital Sant Joan de Déu), University of Barcelona, Barcelona, Spain; Institut d’Investigacions Biomèdiques August Pi i Sunyer (IDIBAPS), Barcelona, Spain; Animal Production Department, Veterinary Faculty, Complutense University of Madrid, Avenida Puerta de Hierro s/n, Madrid 28040, Spain; Institut de Recerca Sant Joan de Déu (IRSJD), Barcelona, Spain; Spanish Network in Maternal, Neonatal, Child and developmental Health Research, RICORS-SAMID, RD24/0013/0004Instituto de Salud Carlos III, Spain; Centre for Biomedical Research on Rare Diseases (CIBER-ER), Madrid, Spain

**Keywords:** Placenta, Metabolism, Pregnancy, Diet, Mediterranean Diet

## Abstract

**Aims:** The Mediterranean diet is associated with reduced cardiometabolic risk, yet its physiological effects during pregnancy and its impact on placental metabolism remain incompletely understood. This study aimed to determine whether maternal adherence to a Mediterranean diet during pregnancy influences placental lipid metabolism and signalling pathways involved in nutrient handling, tissue remodelling, and inflammation, and to assess their relationship with pregnancy outcomes.

**Methods:** Placental samples and clinical outcome data were analysed from pregnant women participating in an unblinded randomized clinical trial of a Mediterranean diet intervention. Placental lipid composition was quantified and the expression of genes and signalling pathways involved in lipid metabolism, nutrient transport, inflammation, and tissue remodelling was evaluated.

**Results:** Maternal adherence to a Mediterranean diet during pregnancy was associated with significant alterations in placental lipid composition, including reduced C18:0 and C24:0 and increased C18:1n9c, C20:3n6, and C22:0, with lower total short-chain fatty acids and higher monounsaturated fatty acids. Placental expression of lipid metabolism regulators ALOX15 and PPARγ was reduced, alongside downregulation of AKT and p38 MAPK signalling pathways. Placentas from mothers adhering to the Mediterranean diet also showed lower expression of amino acid and glucose transporters SLC3A2 and SLC2A1, as well as altered inflammatory and extracellular matrix remodelling markers, including decreased SOCS3 and GHR and increased PAI1 and MMP3.

**Conclusions:** Maternal adherence to a Mediterranean diet during pregnancy modifies placental lipid composition and regulates pathways involved in lipid handling, nutrient transport, inflammation, and tissue remodelling, providing insight into mechanisms linking maternal diet with placental metabolic function.

## 1. Introduction

Diet during pregnancy is a critical determinant of fetal growth^1–3^. Studies in humans and animal models have shown that poor maternal nutrition (both excess and deficiency) from before or during pregnancy can compromise pregnancy outcomes, with both short- and long-term consequences for the offspring ^4–8^. Such consequences include fetal growth restriction, hypoglycemia, and an increased risk of cardiometabolic disorders, including diabetes and hypertension, as well as increased risk of neuropsychiatric conditions, including schizophrenia^9^.

The Mediterranean diet is characterized by a high intake of vegetables, fruits, legumes, nuts, dairy products (principally cheese and yogurt), and extra-virgin olive oil (EVOO); a moderate intake of white meat and fish (particularly oily fish); and limited intake of red and processed meats^10^. It is rich in mono- and polyunsaturated fatty acids and emphasizes foods with antioxidant and anti-inflammatory properties, which protect against oxidative stress, DNA damage, and inflammation while promoting cell proliferation, and angiogenesis^11,12^. Accordingly, adherence to a Mediterranean diet has been associated with a reduced risk of cardiovascular disease, cancer, diabetes, and neurodegenerative disorders ^13–20^.

In pregnant populations, adherence to a Mediterranean diet has also yielded promising outcomes. Several studies have shown that key components of this diet, such as EVOO, nuts, and pistachios, are associated with a reduced risk of gestational diabetes mellitus^21–23^. Recently, a randomised clinical trial (Improving Mothers for a Better PrenAtal Care Trial BarCeloNa, IMPACT BCN) investigated the effects of structured lifestyle interventions, including a Mediterranean diet, in pregnant women at high risk for small-for-gestational-age (SGA) newborns and reported a significant reduction in its prevalence^24^. Moreover, Mediterranean diet intake correlates positively with maternal plasma folate and serum vitamin B12 concentrations^25^, key micronutrients required for DNA synthesis, methylation, and normal fetal development. Indeed, within the same IMPACT BCN trial, offspring of mothers with higher adherence to a Mediterranean diet showed greater total fetal brain volume at magnetic resonance evaluation and higher scores on the Neonatal Neurobehavioral Assessment Scale^26^. The benefits of the Mediterranean diet during pregnancy extend beyond the perinatal period, with improvements in neurodevelopment also observed at two years of age^27^. Similar studies have demonstrated that maternal adherence to this dietary pattern is associated with fetal cardiac structure and function, including increased right ventricular fractional area change and reduced myocardial wall thickness^28^. At the placental level, it has been shown that combining maternal exercise with Mediterranean diet can prevent placental telomere shortening^29^.

Consequently, in recent years, modifying maternal diet to incorporate Mediterranean diet products before or during pregnancy has been considered a promising strategy to prevent or mitigate pregnancy-related complications. Nonetheless, further characterization of the metabolic effects of a Mediterranean diet on placental function, the organ that mediates nutrient and signalling exchange between the mother and fetus, remains to be elucidated. The aim of this study was to investigate the impact of maternal adherence to a Mediterranean diet on placental lipid metabolism and signalling pathways involved in nutrient handling, tissue remodelling, and inflammation, and to examine their association with pregnancy outcomes. To this end, we utilized clinical outcome data and placentas from women with and without adherence to a Mediterranean diet enrolled in the IMPACT BCN trial^24,30,31^.

## 2. Materials and Methods

### 2.1. Participants and Ethics

Placentas were selected from biobank samples collected during the IMPACT BCN trial. This was a parallel, unblinded randomized clinical trial conducted at BCNatal (Hospital Clínic and Hospital Sant Joan de Déu), a large referral center for maternal–fetal and neonatal medicine in Barcelona, Spain. The trial was approved by the Institutional Review Board approval (HCB-2016-0830) and registered at ClinicalTrials.gov (identifier: NCT03166332). Participants were enrolled after eligibility screening during routine second-trimester ultrasound examinations (19.0–23.6 weeks of gestation) and randomized in a 1:1:1 ratio into three groups: a nutritional intervention based on a Mediterranean diet with supplementation of EVOO and walnuts; a Stress reduction program; or a control group without any intervention (usual care). More details are reported in the study protocol elsewhere ^32^.

In all participants of the trial at enrolment (19-23.6 weeks’ gestation) and at final assessment (34-36 weeks’ gestation), a 151-item Food Frequency Questionnaire (FFQ)^33^, and a 17-item Mediterranean Diet Adherence Screener (preg-MEDAS) score, both validated for this study population, were provided^34^. For the present analysis, we considered participants from Mediterranean diet and Usual care groups, non-obese (pre-pregnancy BMI < 30 kg/m²) and with uncomplicated pregnancies, defined as those not affected by SGA, preterm birth, preeclampsia, or gestational diabetes mellitus (n=14 per group). In addition, among women allocated to the Mediterranean diet intervention, only those with high adherence were included, defined as an improvement of ≥3 points in the final preg-MEDAS score^34^.

### 2.2. Mediterranean diet Program

The dietary intervention consisted of monthly individual and group assessments, performed by trained nutritionists, from recruitment (19.0−23.6 weeks’ gestation) until the conclusion of the program (34−36 weeks’ gestation). In each assessment, the goal was to change the overall Mediterranean dietary pattern rather than focusing on single nutrients, including whole-grain cereals (≥5 servings per day); vegetables and dairy products (≥3 servings per day); fresh fruit (≥2 servings per day); and legumes, nuts, fish, and white meat (≥3 servings per week), together with the free provision of EVOO (2 L per month) and walnuts (15 g per day; 450 g per month). Further details are provided in the trial protocol.^32^

### 2.3. Placental sampling

Placental tissue samples were collected at the time of delivery, handled on dry ice to maintain integrity, and stored at −80°C prior to processing. Samples included four blocks containing a full thickness of normal-appearing placental parenchyma (one from each quadrant). After removing membranes, cord, and all blood clots, trimmed placentas were weighed and measured to obtain length, breadth, and thickness. Placental efficiency was calculated as the fetal-to-placental weight ratio.

### 2.4. Placental RNA extraction

Placental RNA was extracted from frozen placental tissues (7 placentas per group and sex, total n=28) using the RNeasy Fibrous Tissue Mini Kit (Qiagen, Hilden, Germany) following the manufacturer’s protocol and as described elsewhere^35^. The quantity of RNA extracted was determined using a NanoDrop spectrophotometer (NanoDrop Technologies, Inc., Auburn, AL), and the RNA was reverse transcribed using a high-capacity cDNA reverse transcription kit (Applied Biosystems, Foster City, USA) according to the manufacturer’s instructions. Quantitative real-time PCR was performed in duplicates using the primer sequences specified in Table S1. The relative mRNA expression levels were normalized to the geometric mean of two AGCCACATCGCTCAGACAC eping genes (*GAPDH* and *HPRT1,* which remained stably expressed across the two groups) and calculated using the 2^−ΔΔCT^ method^36^.

### 2.5. Placental protein extraction

Placental protein extraction was conducted as described previously^37^. Briefly, frozen placental samples were washed in RIPA buffer to remove excess blood, then homogenized using a bead-based method. Following homogenization, samples were incubated in RIPA buffer containing a protease inhibitor cocktail on ice for approximately one hour to allow protein dissociation. Samples were then centrifuged to remove placental debris, and the resulting protein lysates were stored at −80 °C. Total placental protein concentration was determined using the Pierce™ BCA Protein Assay Kit (Thermo Fisher Scientific).

Proteins were separated by electrophoresis and transferred onto 0.2 μm nitrocellulose membranes (Bio-Rad Laboratories Inc., Hercules, CA, USA). Membranes were blocked with fetal bovine serum and/or skimmed milk, depending on whether phosphorylated or total proteins were analyzed, respectively, and incubated overnight at 4 °C with primary antibodies (Table S2). The following day, membranes were washed several times with TBS-Tween and incubated with HRP-conjugated secondary antibodies (1:10,000 dilution; NA931 or NA934; Amersham ECL Mouse/Rabbit IgG). Immunocomplexes were detected using SuperSignal™ West Femto Maximum Sensitivity Substrate (Thermo Fisher Scientific) and imaged with the iBright 1500 Imaging System. To control for variations in protein loading, membranes were stained with Ponceau S, and signal intensities were normalized to Ponceau S staining or to total protein levels, as appropriate.

### 2.6. Placental lipid analysis

For lipidomics analysis, placental samples were converted into fatty acid methyl esters (FAMEs) using the method described elsewhere^38,39^. Briefly, FAMEs were derived from freeze-dried placentas by employing a solution of 10% acetyl chloride in methanol and toluene, heated at 70°C for 120 minutes. Post-heating, 6% potassium carbonate and toluene were introduced, followed by centrifugation at 1,500g for 5 minutes. The resulting organic layers containing FAMEs were segregated into individual vials and stored at −20°C for subsequent analysis.

The FAMEs were analyzed using a gas chromatograph (Varian CP-3800; Agilent, USA) equipped with a flame ionization detector and an SP-2560 column (Supelco, Bellefonte, PA, USA). The analytical process involved injecting 1.0 μL of the sample in split mode at a 1:30 split ratio. The detector and injector oven temperatures were maintained at 260°C. The oven temperature profile began at 140°C for 5 minutes, increased by 4°C per minute to 240°C, then increased by 20°C within 1 minute to 260°C, where it remained constant for 15 minutes. Identification of FAMEs was accomplished by comparison with a standard FAME mixture (Supelco® 37-component FAME Mix; Sigma Aldrich). Results were presented as a percentage of the total identified FAMEs.

### 2.7. Statistical analysis

Maternal data were analyzed using Student’s *t*-test or Chi square test. Newborn weight and placental morphometric parameters, as well as all placental lipid profiling and gene expression data, were analyzed by two-way ANOVA (diet and newborn sex), followed by Tukey’s multiple-comparisons test. Assumptions of normality and homogeneity of variances were verified using the Shapiro–Wilk test and Bartlett’s test, respectively. Western blot data were analysed by Student’s *t*-test for each newborn sex separately. A *P* value < 0.05 was considered statistically significant.

## 3. Results

### 3.1. Maternal Characteristics, newborn weights and placental macroscopic phenotype

No differences were observed in maternal baseline characteristics or perinatal results among the study groups (Table 1). In line with the main results from the IMPACT BCN trial, a tendency to lower SGA incidence was observed in the intervention groups^40^.

**Table 1.** Maternal baseline and perinatal characteristics. Data are given as mean (SD) or n (%). BMI, body mass index; †Socioeconomic status is defined as follows: low (never worked or unemployed > 2 years) medium (secondary studies and work) and high (university studies and work).

|  | **Mediterranean Diet** | **Usual care** | **Mediterranean Diet vs. Usual care** |
| --- | --- | --- | --- |
|  | n = 14 | n = 14 | *p* value |
| **Maternal baseline characteristics** |  |  |  |
| **Age (years)** | 36.5 (4.3) | 34.9 (7.3) | 0.47 |
| **Ethnicity** |  |  | 0.28 |
| Afro-American | 0 (0) | 0 (0) |  |
| Asian | 0 (0) | 0 (0) |  |
| Latin | 3 (21.4) | 1 (7.1) |  |
| White | 11 (78.6) | 13 (92.9) |  |
| **Study class** |  |  | 0.38 |
| No education/Primary grade | 0 (0) | 1 (7.1) |  |
| Secondary/Technology grade | 5 (35.7) | 7 (50.0) |  |
| University | 9 (64.3) | 6 (42.9) |  |
| **Socio-economic status†** |  |  | 0.38 |
| Low | 0 (0) | 1 (7.1) |  |
| Medium | 5 (35.7) | 7 (50.0) |  |
| High | 9 (64.3) | 6 (42.9) |  |
| **BMI before pregnancy (Kg/m<sup>2</sup>)** | 22.8 (2.5) | 23.4 (3) | 0.55 |
| **Medical condition** |  |  |  |
| Autoimmune disease | 1 (7.1) | 3 (21.4) | 0.28 |
| Chronic hypertension | 1 (7.1) | 0 (0) | 0.31 |
| Psychiatric disorders | 1 (7.1) | 2 (14.3) | 0.54 |
| **Nulliparous** | 10 (71.3) | 8 (57.1) | 0.43 |
| **Assisted reproductive technologies** | 0 (0) | 3 (21.4) | 0.07 |
| **During pregnancy** |  |  |  |
| Cigarette smoking | 1 (7.1) | 4 (28.6) | 0.14 |
| Alcohol intake | 0 (0) | 0 (0) | 1.00 |
| Drugs consumption | 1 (7.1) | 0 (0) | 0.31 |
| Gestational age at recruitment (weeks) | 20.7 (0.5) | 20.8 (0.6) | 0.59 |
| **Perinatal data** |  |  |  |
| Gestational age at delivery (weeks) | 40 (0.9) | 39.8 (1.2) | 0.49 |
| Caesarean section | 2 (14.3) | 5 (35.7) | 0.19 |

Regarding newborn and placental macroscopic phenotypes, we observed no differences in birth weight, placental weight, placental efficiency), or placental length, breadth, or thickness between women in the usual care and Mediterranean diet groups (Figure 1A-F).

**Figure 1.**
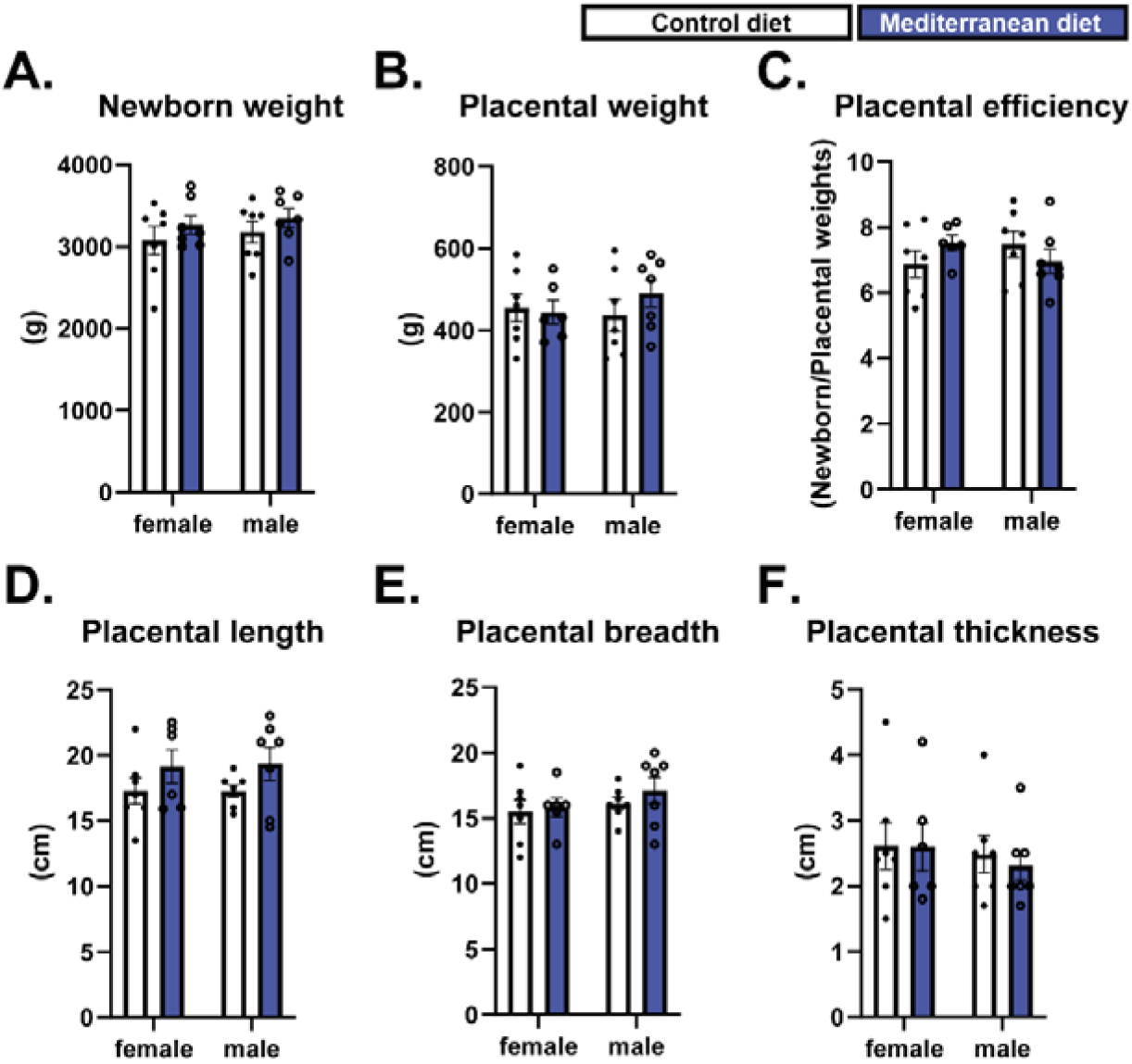
Newborn biometric parameters and placental characteristics. Newborn biometric data, placental weight, placental efficiency (defined as the ratio of fetal to placental weight), and placental macroscopic parameters. Data were analysed by two-way ANOVA followed by Tukey’s post hoc test. Data are shown as individual values, with columns representing the mean ± SEM.

### 3.2. Mediterranean diet induced changes in placental fatty-acid lipid species

The total content of 16 lipid species within the placenta was ascertained by lipidomic analysis. Principal component analysis showed modest clustering of placentas from usual care and Mediterranean diet groups, indicating diet-associated variation in placental lipid composition (Fig. 2A). To identify lipid species contributing most strongly to dietary discrimination, we performed a supervised Random Forest analysis (Fig. 2B). Among the highest-ranking lipids were the long-chain saturated fatty acid C24:0, total saturated fatty acid (SFA), and C18:0, suggesting that saturated lipid species were major contributors to the separation between dietary groups.

**Figure 2.**
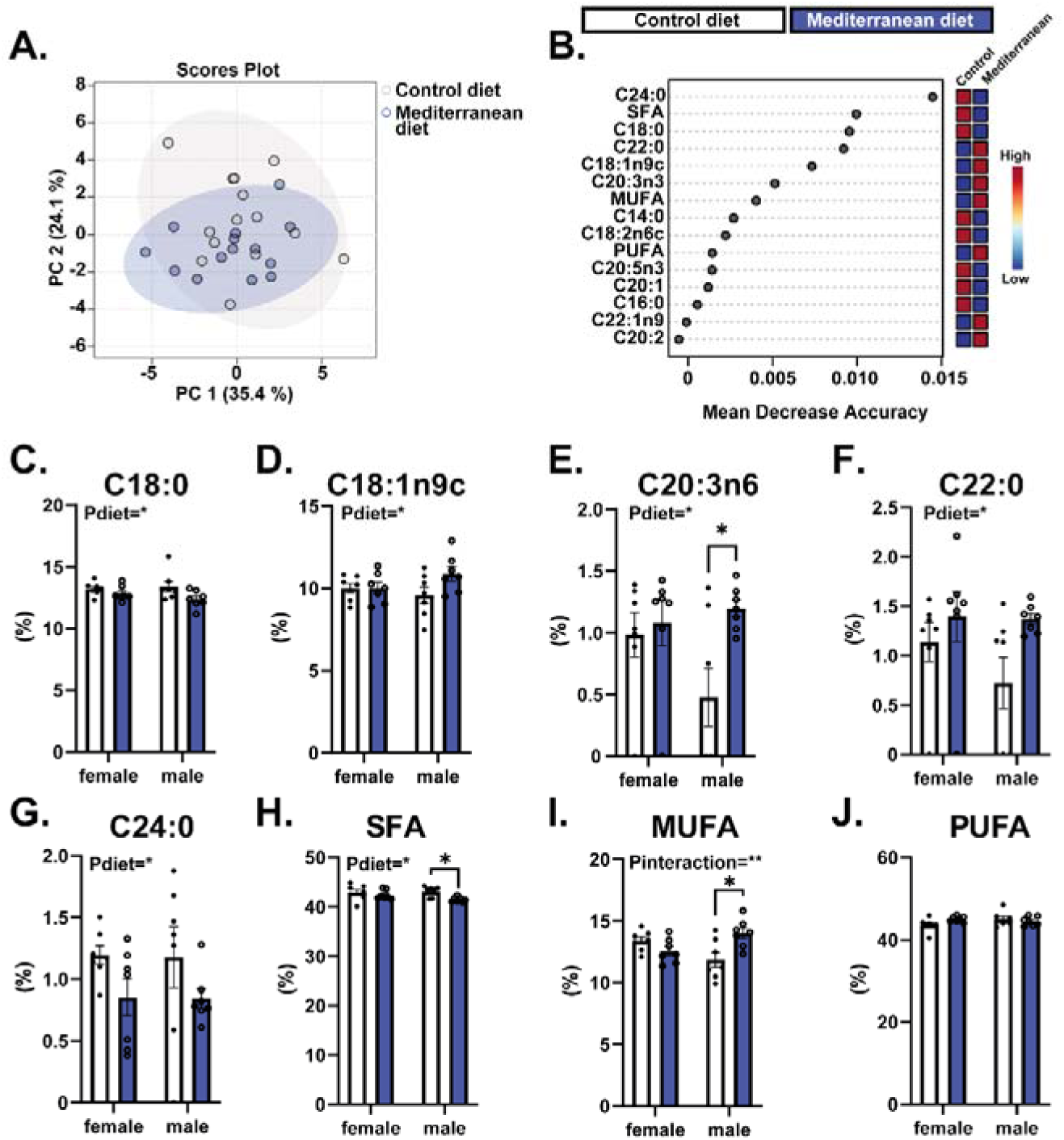
Mediterranean diet alters placental lipid composition. (A,B) Principal component analysis (PCA) and random forest analysis of placental lipid profiles. (C–G) Placental lipid species significantly altered by maternal adherence to the Mediterranean diet. Lipid species are expressed as the proportion (%) of total measured placental lipids. Saturated fatty acids (SFA) were defined as the sum of C14:0, C16:0, C18:0, C20:0, C22:0 and C24:0; monounsaturated fatty acids (MUFA) as the sum of C16:1, C18:1n9c, C20:1 and C22:1n9; and polyunsaturated fatty acids (PUFA) as the sum of C18:2n6c, C20:2, C20:3n6, C20:3n3, C20:4n6, C20:5n3 and C22:6n3. Data were analysed by two-way ANOVA followed by Tukey’s post hoc test. *P<0.05. Data are shown as individual values, with columns representing the mean ± SEM.

Of the 16 lipid species quantified (Fig.2 and Fig. SX), five were significantly altered by a maternal Mediterranean diet. Specifically, placentas from the Mediterranean diet group exhibited reduced levels of the saturated fatty acids C18:0 and C24:0 (Fig. 2C, G). In contrast, the monounsaturated fatty acid (MUFA) C18:1n9c and the polyunsaturated fatty acid (PUFA) C20:3n6, as well as the long-chain saturated fatty acid C22:0, were significantly increased in placentas exposed to the Mediterranean diet (Fig. 2D–F). Notably, the increase in C20:3n6 was particularly pronounced in male placentas, indicating a sex-specific dietary effect.

When lipid species were grouped by class, maternal adherence to a Mediterranean diet was associated with a significant reduction in total saturated fatty acid (SFA), driven predominantly by changes in male placentas (Fig. 2H). Conversely, total MUFA were significantly elevated in male placentas (Fig. 2I), whereas no significant differences were observed in total PUFA levels (Fig. 2J). Taken together, these data suggest that a Mediterranean diet affects the placental lipid profile.

### 3.3 Mediterranean diet alters select cellular signalling pathways in the placenta

Exposure to fatty acids can alter the phosphorylation status of key metabolic signalling pathways, including AKT and MAPKs^41–43^. In Mediterranean diet-exposed placentas, we observed a significant reduction in phosphorylated AKT levels in male, but not female, placentas, without changes in total AKT protein abundance (Fig. 3A–C). Moreover, the phosphorylated levels of p38 MAPK, a signalling pathway involved in multiple other cellular processes including vascular growth^44–46^, were significantly reduced in both male and female placentas. Notably, total p38 MAPK protein levels were significantly increased in male placentas (Fig. 3D–F). Other key components of other metabolic and growth-related pathways, namely PI3K and mTOR (PI3K-p85α, mTOR-GbL, and RAPTOR), were not changed by a maternal Mediterranean diet (Figure 3). Taken together, these data suggest that a maternal Mediterranean diet modulates select metabolic and growth signalling pathways in the human placenta.

**Figure 3.**
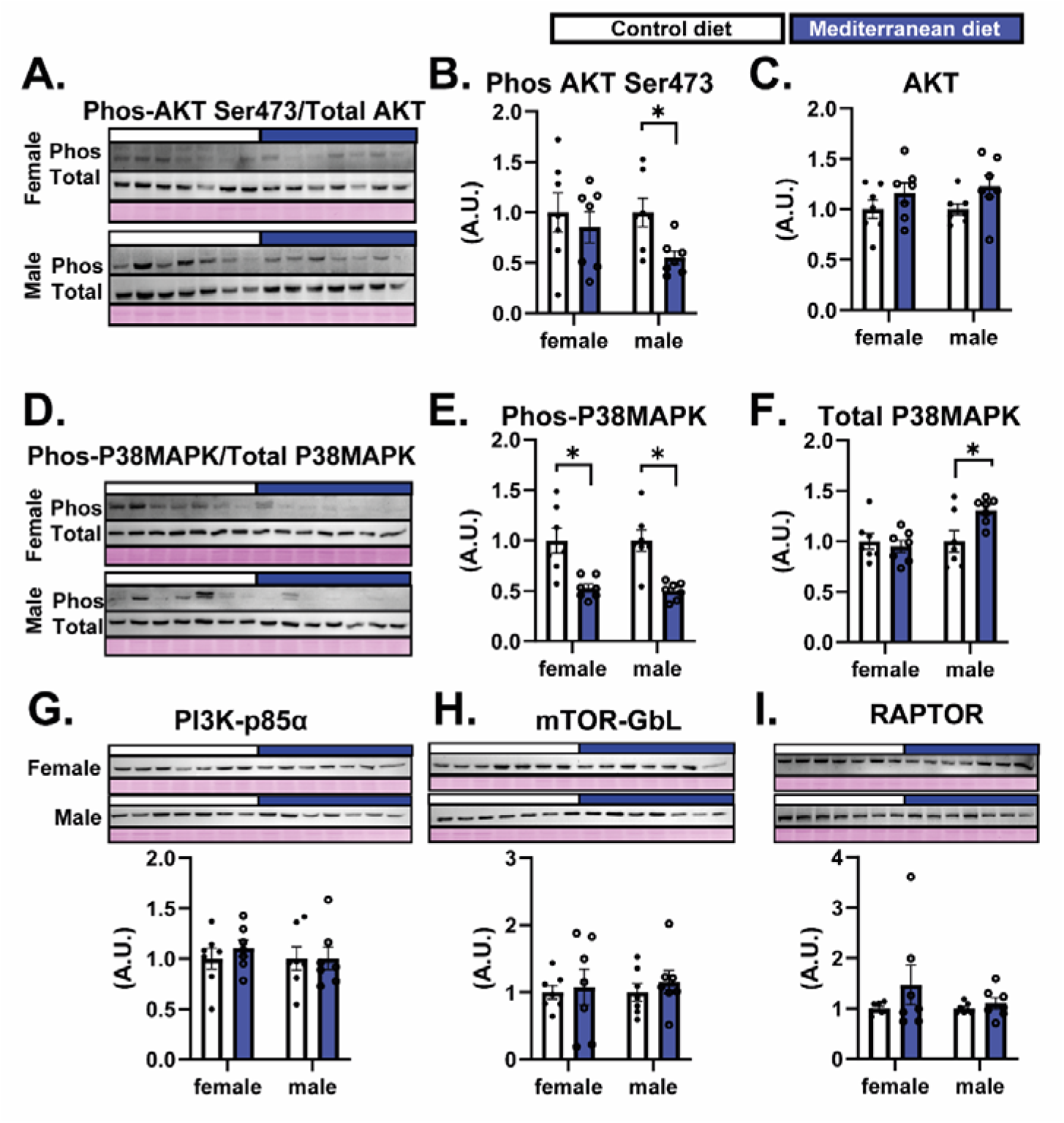
Mediterranean diet alters placental signaling pathways. Data were analyzed separately using Student’s *t*-test. \**P* < 0.05 was considered statistically significant. Data are shown as individual values, with columns representing the mean ± SEM. Levels of phosphorylated proteins were normalized to the corresponding total protein abundance, and total protein levels were normalized to Ponceau S staining.

### 3.3 Mediterranean diet reduces placental expression of nutrient transporters and lipid metabolism–related and regulatory genes

We next analysed the mRNA expression of three key nutrient transporters: *SLC3A2*, which is involved in amino acid transport and trophoblast function^47^; *SLC2A1*, a major glucose transporter^48^; and *SLC2A8*, a class III facilitative glucose/fructose transporter^49^. This revealed that a maternal Mediterranean diet significantly downregulated *SLC3A2* and *SLC2A1* mRNA expression, whereas *SLC2A8* expression remained unchanged (Fig. 4A-C).

**Figure 4.**
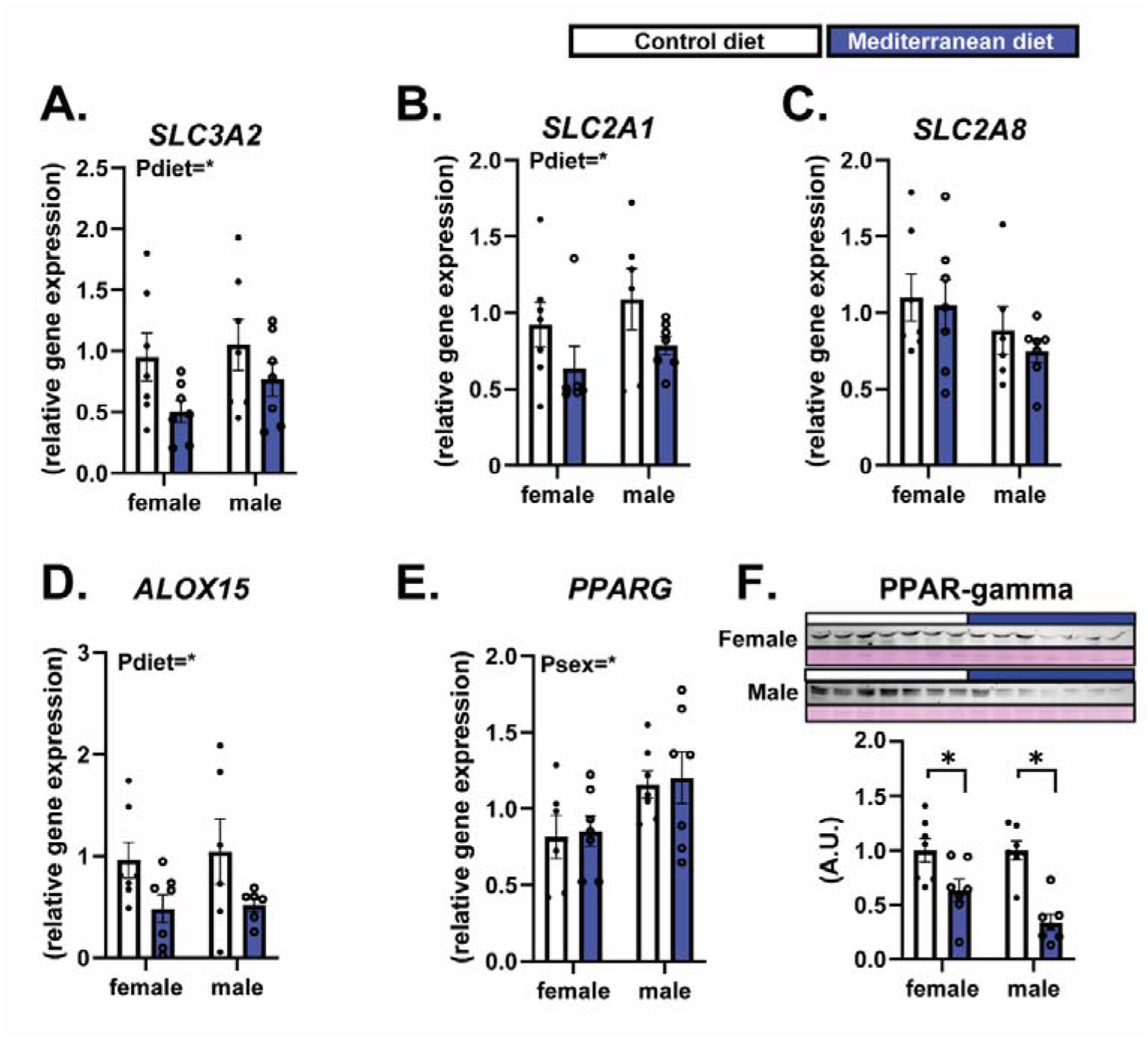
Mediterranean diet reduces mRNA levels of key nutrient transporters and decreases the abundance of lipid-handling genes and proteins (ALOX15 and PPARγ). Gene expression data were analyzed by two-way ANOVA followed by Tukey’s post hoc test. PPARγ protein levels were analyzed using Student’s *t*-test and normalized to Ponceau S staining. \**P* < 0.05 was considered statistically significant. Data are shown as individual values, with columns representing the mean ± SEM.

We also measured the mRNA levels of genes involved in mitochondrial fatty acid β-oxidation (*ACADL*, *ACADM*, *ACADVL*), fatty acid synthesis (*FASN*), and lipid transport (*APOE*). However, none of these genes were altered by a maternal Mediterranean diet (Fig. S2A-E). In contrast, *ALOX15,* a gene involved in lipid peroxidation, was significantly reduced in placentas from the Mediterranean diet group (Fig. 4D). Furthermore, although mRNA levels were unchanged, the abundance of PPARG, a conserved regulator of placentation^50^ and PUFA metabolism, was significantly reduced in both male and female placentas from the Mediterranean diet group (Fig. 4E-F). Taken together, these findings indicate that Mediterranean diet selectively modulates nutrient transport and lipid peroxidation pathways in the placenta.

### 3.4 Mediterranean diet modulates placental inflammation and extracellular matrix remodelling genes and proteins

Inflammation and extracellular matrix (ECM) remodelling are tightly coupled processes, as inflammatory signalling induces ECM degradation and reorganization through the regulation of proteases and matrix components, while changes in ECM composition and structure feedback to modulate inflammatory responses and tissue function. In light of this interplay, and given that consumption of Mediterranean diet components has been linked to reduced inflammation^51^, we next examined the expression of genes involved in inflammatory signalling and ECM remodelling. Proteins encoded by the *RAGE*, *PTGS2*, and *SOCS3* genes are known to coordinate cytokine-driven responses^52–54^, while those encoding *GHR*, *PAI1*, and *MMP3* contribute to ECM remodelling^55–57^. While no differences were observed in the mRNA levels of *RAGE* or *PTGS2*, *SOCS3* expression was significantly reduced in female placentas of mothers adhering to the Mediterranean diet (Fig. 5A-C). In contrast, we found a significant overall reduction in the *GHR* mRNA levels in the placentas of the Mediterranean diet group (Figure 5D). At the protein level, PAI1 and MMP3 were significantly increased in female placentas from the Mediterranean diet group, whereas in males, only PAI1 abundance was elevated, with no change observed for MMP3 (Fig. 5E-F). Taken together, these data indicate that Mediterranean diet modulates the expression of genes and the abundance of proteins involved in inflammation and extracellular matrix remodelling in the placenta.

**Figure 5.**
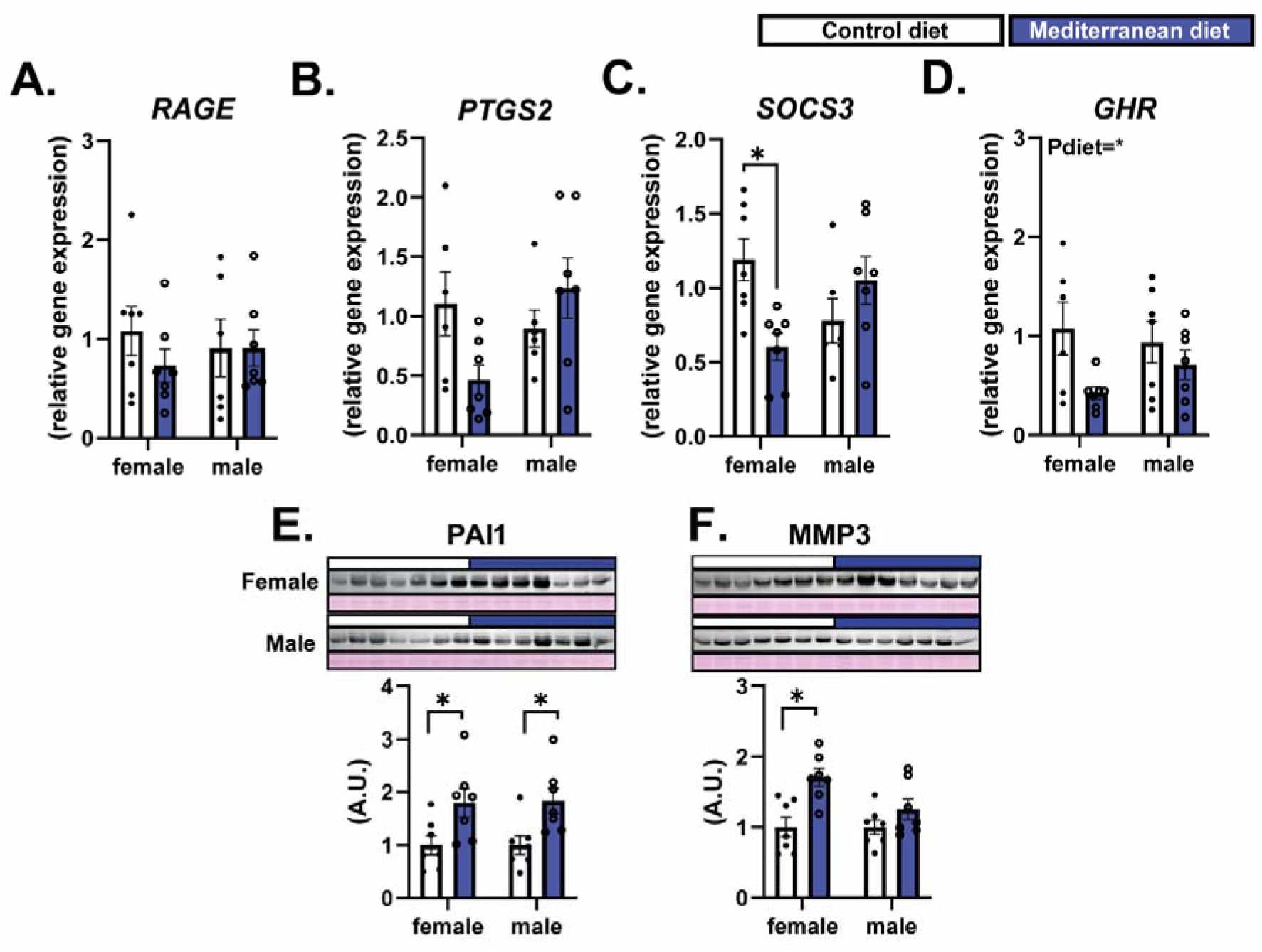
Mediterranean diet reduces mRNA levels of genes involved in placental inflammation and growth signaling, while increasing proteins associated with extracellular matrix remodeling. (A–D) Relative mRNA expression of RAGE, PTGS2, SOCS3, and GHR in placental tissue. (E–F) Protein abundance of PAI1 and MMP3 with corresponding immunoblots. Protein levels were normalized to Ponceau S staining. Gene expression data were analyzed by two-way ANOVA followed by Tukey’s post hoc test. Protein data were analyzed using Student’s t-test. P < 0.05 was considered statistically significant. Data are shown as individual values, with columns representing the mean ± SEM.

## 4. Discussion

In this study, we show that maternal adherence to a Mediterranean diet alters placental lipid composition and modulates signalling pathways, nutrient transporter expression, and inflammatory and extracellular matrix–related pathways. Several of these effects were sex-specific, suggesting differential responses of male and female placentas to maternal dietary exposure. Notably, these molecular changes occurred in the absence of differences in placental macroscopic phenotype or neonatal weight. These findings provide insight into the mechanisms through which maternal diet may influence placental function.

Consistent with these observations, we observed significant alterations in five placental lipid species, along with key genes and proteins involved in lipid handling, including *ALOX15*^58^ and its downstream target, PPAR gamma^59^ (both reduced in the Mediterranean diet group). Among these, the saturated long-chain fatty acid C18:0 (stearic acid) was significantly reduced in placentas from women consuming a Mediterranean diet. This finding is of interest given that accumulation of saturated fatty acids has been shown to activate pro-inflammatory pathways, including increased cytokine production^60^. In line with previous reports linking elevated C18:0 to metabolic dysfunction, women with GDM exhibit higher circulating levels of C18:0 in maternal plasma^61^, and studies in primary human trophoblasts have demonstrated that C18:0 stimulates the synthesis and release of pro-inflammatory cytokines, including TNF-α, IL-6, and IL-8^62^.

In addition, the proportion of C24:0 (lignoceric acid) was significantly reduced relative to other detected lipid species in the placenta of the Mediterranean diet group. Although C24:0 is present at low abundance in the placenta, its levels are responsive to maternal dietary exposures, as demonstrated in animal models subjected to obesogenic diets during gestation^63^. Studies in both human cohorts and animal models have identified C24:0 as a lipid species enriched in obese and diabetic states, and fetal programming studies using high-energy diets during gestation have shown that such exposures can induce adipose tissue dysfunction in the offspring^64,65^. Accumulation of C18:0 and C24:0 has been associated with mitochondrial dysfunction, characterised by reduced oxidative capacity, increased fatty acid accumulation, and insulin resistance^64,65^. Although these observations require further validation in placental tissue, the reduced abundance of C18:0 and C24:0 may contribute to improved mitochondrial oxidative capacity within the placenta, potentially limiting lipid accumulation and supporting insulin sensitivity and metabolic efficiency. Even in the absence of differences in birthweight, such placental adaptations could influence the intrauterine environment and may have implications for long-term offspring health.

Among the lipid species increased in placentas from the Mediterranean diet group, C22:0 (behenic acid) warrants particular attention. Studies in animal models have shown that C22:0 can alleviate inflammation and insulin resistance in GDM, at least in part by modulating the TLR4/NF-κB signalling pathway. In addition, *in vitro* studies using placental tissues have demonstrated that this fatty acid reduces the secretion of pro-inflammatory cytokines, including IL-6, IL-17, and TNF-α, as well as chemokines such as CCL3, CCL8, CXCL2, and CXCL^66^.

We also observed a sexually dimorphic response in the proportion of C20:3n-6 (dihomo-γ-linolenic acid), a precursor for eicosanoid synthesis, which was significantly increased in male but not female placentas in the Mediterranean diet group. In the control group, male placentas, on average, had lower levels of this fatty acid than female placentas, but this difference was not statistically significant. Moreover, information regarding the role of this lipid species in placental function remains limited. Future studies are needed to determine whether changes in placental lipid species handling with the maternal Mediterranean diet reflect increased peroxisomal fatty acid oxidation, a shift away from lipid storage pathways toward enhanced lipid catabolism in the placenta, or increased transplacental transfer of these fatty acids to the developing fetus.

Both AKT and p38MAPK are critical signalling pathways important for metabolism, inflammation, and cell growth ^67–69^. Phosphorylation of both pathways was significantly reduced in the placentas of mothers following a Mediterranean diet. Notably, these changes occurred in the absence of alterations in the upstream regulatory unit of PI3K-p85 or in the downstream mediators growth and nutrient-sensing mediators, mTOR, and RAPTOR. Although further investigation is required, these findings suggest that the observed changes may reflect alterations in specific input signals, such as lipid species, membrane-associated receptors, or hormonal cues, rather than changes in the expression of core components of the PI3K–mTOR signalling axis.

In support of this interpretation, p38MAPK activity has been shown to promote the expression of suppressor of cytokine signalling 3 (SOCS3)^70^, and we observed significantly reduced SOCS3 mRNA levels in female placentas exposed to the Mediterranean diet. Moreover, studies in non-placental systems have demonstrated that growth hormone receptor (GHR) deficiency inhibits activation of the PI3K–AKT signalling pathway^71^. Consistent with this, we detected an overall reduction in GHR mRNA levels in the Mediterranean diet group, which may contribute to the attenuation of AKT signalling observed in these placentas. In addition, we observed a significant reduction in SLC3A2 expression in the Mediterranean diet placentas. SLC3A2 is an integrin-associated protein that forms the heavy chain of heteromeric amino-acid transporters and modulates integrin-dependent signalling in the placenta. Knockdown of SLC3A2 has been shown to reduce AKT phosphorylation^72,73^, suggesting that decreased *SLC3A2* expression may further contribute to reduced AKT activity. Notably, we also detected reduced mRNA levels of *SLC2A1 (GLUT1)*, the primary placental glucose transporter. While *SLC2A1* expression is regulated by multiple signalling and metabolic cues, including growth factor and insulin-related pathways^74,75^, its reduction may reflect broader alterations in nutrient-sensing and transport mechanisms associated with attenuated AKT signalling, although a direct causal relationship cannot be inferred from the present data and will require further work.

Finally, we detected increased protein levels of plasminogen activator inhibitor-1 (PAI1; SERPINE1) in both male and female placentas exposed to the Mediterranean diet. Although elevated PAI1 levels have been associated with pregnancy complications such as GDM, preeclampsia, and fetal growth restriction^56^, PAI1 also plays an essential physiological role in placental development. This protein is expressed in extravillous interstitial and vascular trophoblasts, where it contributes to the regulation of extracellular matrix degradation, thereby controlling trophoblast invasion and ensuring appropriate remodelling of maternal spiral arteries while preventing excessive invasion^56^. Interestingly, in female placentas exposed to the Mediterranean diet, we also observed increased levels of matrix metalloproteinase-3 (MMP3). Under physiological conditions, PAI1 participates in the fine-tuning of matrix metalloproteinase activity by regulating the plasminogen–plasmin system. The concomitant increase in PAI1 and MMP3 in female placentas may therefore reflect a balanced remodelling programme rather than a pathological phenotype, although further studies will be required to define the functional consequences of this coordinated regulation.

Importantly, despite the observed molecular and biochemical adaptations observed in the placenta, neither newborn birth weight nor gross placental phenotype differed between the Mediterranean diet and control groups, irrespective of fetal sex. These findings suggest that maternal adherence to the Mediterranean diet may promote subtle functional and molecular placental adaptations without altering overall fetal growth at birth. However, whether these placental changes confer long-term benefits to the offspring remains unknown. Longitudinal follow-up studies will be essential to determine whether such adaptations influence susceptibility to metabolic, neurological, or cardiovascular disorders later in life.

In conclusion, this study provides evidence that maternal adherence to the Mediterranean diet modulates placental lipid composition, nutrient transport, and signalling pathways controlling growth and inflammation, highlighting the placenta as a key mediator of maternal diet-fetal interactions. These findings support the Mediterranean diet as a promising, non-pharmacological strategy to improve placental metabolic function during pregnancy. However, larger and well-powered studies integrating state-of-the-art approaches, including transcriptomics, proteomics, metabolomics, and detailed histological analyses, will be required to fully characterize the impact of maternal dietary patterns on placental phenotype and to assess the long-term benefits of these changes for maternal and offspring health.

## Data availability

The datasets are available from the corresponding author upon reasonable request.

## Author Contributions

acquired human placental samples (LY, AC, FC, EG, FC), lipid analysis (RB-P, AC, JDLF), laboratory work (JL-T), acquired funding (JL-T, ANS-P, LY, AC, FC, EG, FC), prepared the figures (JL-T), designed laboratory experiments (JL-T, ANS-P), data analysis (JL-T, LY, RB-P), drafted original version of the article (JL-T), all authors proofread the final version of the manuscript.

## Acknowledgment

We thank the Clinic-IDIBAPS Biobank and the “Biobanc de l’Hospital Infantil Sant Joan de Déu per a la Investigació” integrated in the Spanish Biobank Network of ISCIII for the sample and data procurement.

---

**Figure S1.**
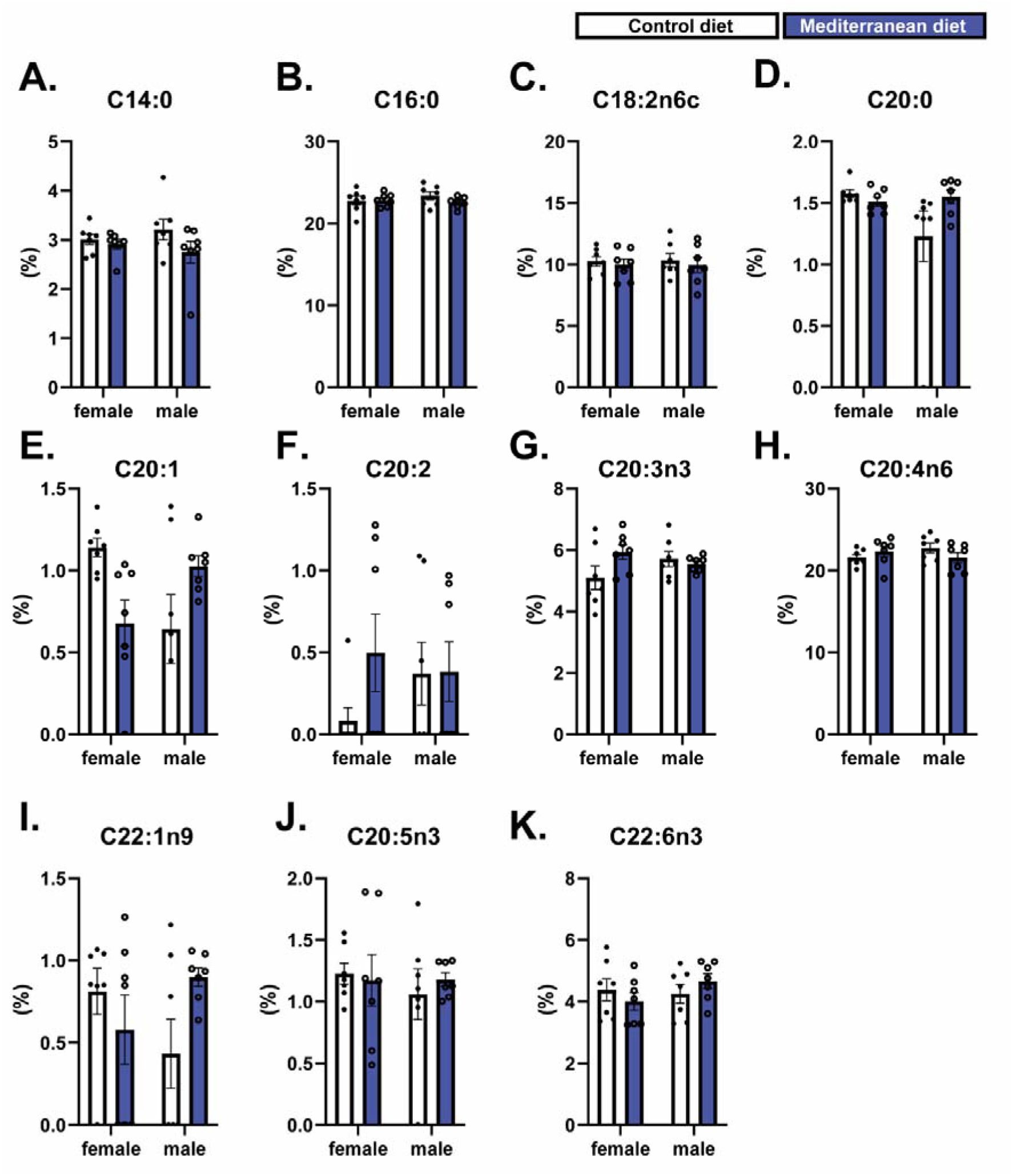
Additional list of lipids measured in placental samples. Data were analysed by two-way ANOVA followed by Tukey’s post hoc test. Data are shown as individual values, with columns representing the mean ± SEM.

**Figure S2.**
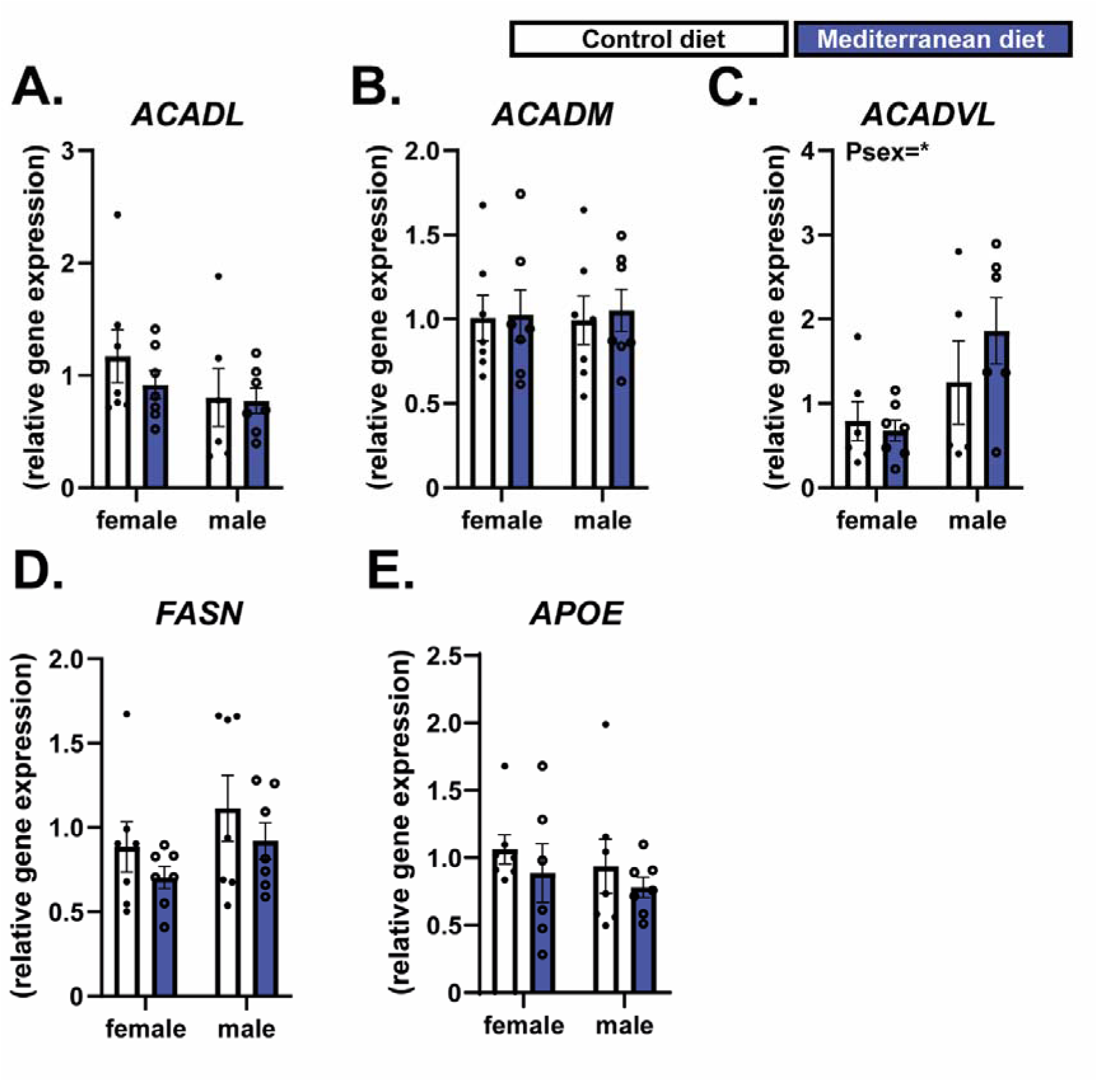
Additional genes involved in lipid handling measured in placental samples. Data were analysed by two-way ANOVA followed by Tukey’s post hoc test. Data are shown as individual values, with columns representing the mean ± SEM.

**Table S1.**
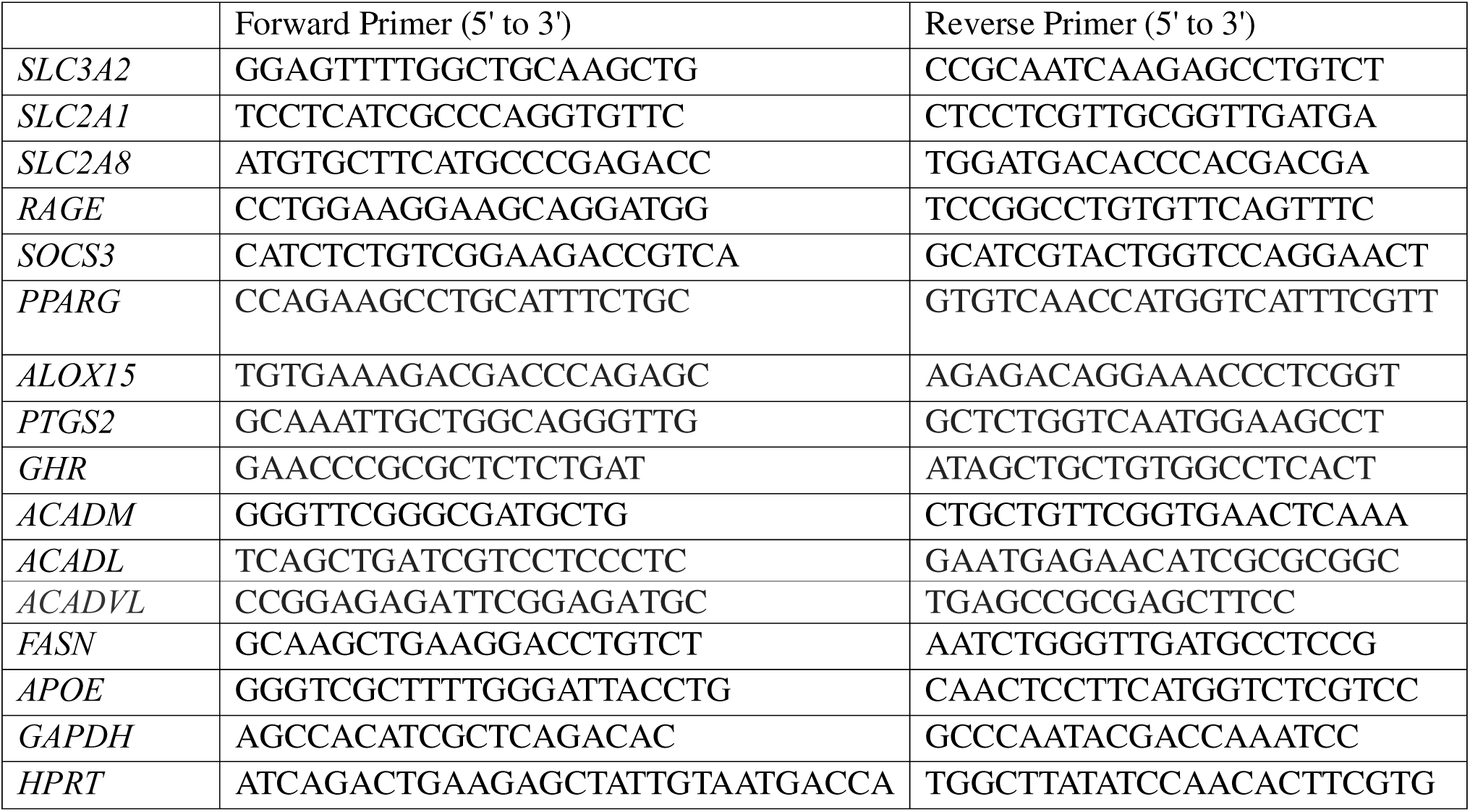
List of primers used for qPCR.

**Table S2.**
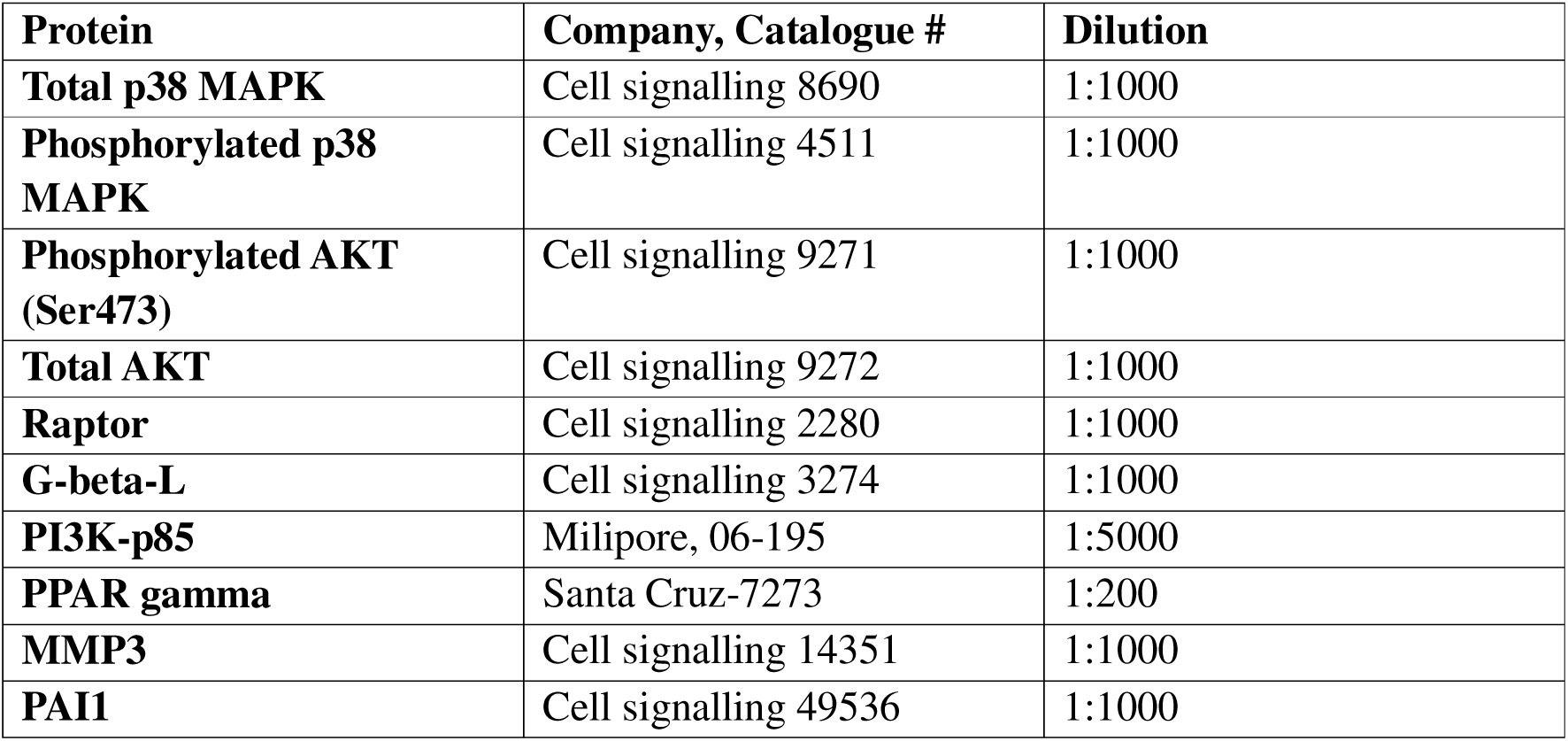
List of antibodies used for western blotting.

| Protein | Company, Catalogue # | Dilution |
| --- | --- | --- |
| Total p38 MAPK | Cell signalling 8690 | 1:1000 |
| Phosphorylated p38 MAPK | Cell signalling 4511 | 1:1000 |
| Phosphorylated AKT (Ser473) | Cell signalling 9271 | 1:1000 |
| Total AKT | Cell signalling 9272 | 1:1000 |
| Raptor | Cell signalling 2280 | 1:1000 |
| G-beta-L | Cell signalling 3274 | 1:1000 |
| PI3K-p85 | Milipore, 06-195 | 1:5000 |
| PPAR gamma | Santa Cruz-7273 | 1:200 |
| MMP3 | Cell signalling 14351 | 1:1000 |
| PAI1 | Cell signalling 49536 | 1:1000 |

